# Reliability of brain metrics derived from a Time-Domain Functional Near-Infrared Spectroscopy System

**DOI:** 10.1101/2024.03.12.584660

**Authors:** Julien Dubois, Ryan M. Field, Sami Jawhar, Erin M. Koch, Zahra M. Aghajan, Naomi Miller, Katherine L. Perdue, Moriah Taylor

## Abstract

With the growing interest in establishing brain-based biomarkers for precision medicine, there is a need for noninvasive, scalable neuroimaging devices that yield valid and reliable metrics. Kernel’s second-generation Flow2 Time-Domain Functional Near-Infrared Spectroscopy (TD-fNIRS) system meets the requirements of noninvasive and scalable neuroimaging, and uses a validated modality to measure brain function. In this work, we investigate the test-retest reliability (TRR) of a set of metrics derived from the Flow2 recordings. We adopted a repeated-measures design with 49 healthy participants, and quantified TRR over multiple time points and different headsets—in different experimental conditions including a resting state, a sensory, and a cognitive task. Results demonstrated high reliability in resting state features including hemoglobin concentrations, head tissue light attenuation, amplitude of low frequency fluctuations, and functional connectivity. Additionally, passive auditory and Go/No-Go inhibitory control tasks each exhibited similar activation patterns across days. Notably, areas with the highest reliability were in auditory regions during the auditory task, and right prefrontal regions during the Go/No-Go task, consistent with prior literature. This study underscores the reliability of Flow2-derived metrics, supporting its potential to actualize the vision of using brain-based biomarkers for diagnosis, treatment selection and treatment monitoring of neuropsychiatric and neurocognitive disorders.

## Introduction

Test-retest reliability (TRR) of non-invasive functional neuroimaging-derived metrics has become an increasingly important avenue of research as promising evidence for brain-based biomarkers and clinical use cases has grown over recent years. In order for a metric to have therapeutic or diagnostic utility, it must be stable in the absence of structural or functional changes in an individual. Under this condition, any change in the metric can be attributed to a bona fide underlying change in the brain-behavior axis, which may reflect disease progression or response to clinical intervention^1,2^.

The strictest form of TRR consists in repeating a measurement with the same device, on the same day, in the same individual, assuming nothing changed between the two assessments. However, the reliability of interest for brain-based biomarkers is of a broader form^3^, requiring stability across devices and days. Above all, measurement variability within an individual (which may stem from different confounding factors, such as time of day, exercise, sleep, neuroactive substances) should be small in relation to between-subject variability (for diagnostic purposes) and to within-subject variability across functional changes of interest (for treatment outcome purposes).

Furthermore, reliability alone isn’t sufficient for clinical utility. A measurement must also be valid, i.e. pertain to the function that it intends to measure. Indeed, a very reliable measure could be a reflection of a stable confound which is irrelevant to the function of interest. On the other hand, an observed lack of reliability does not readily invalidate a measure; it may simply indicate that the function of interest was not properly held constant across measurements, or may represent a valid change. While the bulk of non-invasive functional neuroimaging studies has sought to validate derived neural metrics, a growing body of literature has explored the TRR of these measurements in neuropsychiatric conditions such as depression^1,2^, cognitive decline^4,5^, pain^6^, stroke^7^, aging^8^, and ADHD^9^. These studies are helpful to establish the current standard of TRR across common noninvasive neuroimaging modalities (functional Magnetic Resonance Imaging, fMRI; electroencephalography, EEG; and functional Near-Infrared Spectroscopy, fNIRS).

At the forefront of noninvasive neuroimaging is Functional Magnetic Resonance Imaging (fMRI), which has progressed our understanding of the human brain both in healthy and clinical populations. However, the within-subject stability of fMRI recordings has been called into question, and the literature surrounding TRR, while extensive, is nuanced at best. These findings may not be surprising given the inherent challenges that fMRI faces in collecting homogeneous data, such as motion, system artifacts, and variability across different scanners and sites^3^. On one hand, there have been resting-state studies that demonstrate good to excellent reliability (particularly with within-visit repetition) of extracted neural features in some brain regions or when considering specific functional brain networks^10,11^. The details of these papers, nonetheless, shed some light on the incongruence of the literature (e.g., they showed altered reliability with different processing steps). On the other hand, meta-analyses have found standard measures of reliability to range from poor to fair, across a variety of tasks^12,13^. In the face of this, numerous studies have proposed best practices for enhancing and interpreting fMRI reliability^14,15^, emphasizing that it is best done alongside the assessment of validity.

While fMRI measures hemodynamic signals, electroencephalography (EEG) measures electrical brain activity. The TRR of both time- and frequency-domain EEG metrics has been investigated. The literature suggests that power spectra from resting state measurements are more reliable than task-evoked event related potentials (ERPs)^16^. For resting state, the measured reliability varies across frequency bands (e.g., alpha vs. others) and spatial location of the electrodes (e.g. central vs. peripheral)^16^. For tasks, it has been shown that ERP reliability is affected by task type^17^, varies across different ERP components (e.g. P3 vs. N2)^17,18^, and across the ERP time course^18^. Akin to differences in fMRI scanners across different locations, EEG variability could stem from different operators, the use of different caps, and different placements of the same caps^19–21^. Taken together, the clinical utility of EEG-derived metrics may be limited to those measures with high reliability (e.g., certain frequency bands) and their corresponding experimental designs (e.g., resting state). Another viable modality with fast recording time scales is Magnetoencephalography (MEG). However, metrics derived from MEG are not considered here, as its cost and ecological validity are far from that needed for precision medicine.

In recent years, Functional Near-Infrared Spectroscopy (fNIRS) has become an increasingly attractive neuroimaging modality, as it is portable, affordable, and robust to motion artifacts^22^. This method, which uses light to measure brain function, can be particularly suitable for measuring clinical populations as it is participant-friendly—there are no loud scanner noises, confinement concerns, or messy gel to apply to the scalp to make good contact. There are also minimal restrictions on movement and speech^23,24^. In this domain, there have been a number of promising studies exhibiting good to excellent reliability across a multitude of tasks including resting state^25,26^, visual and auditory sensory tasks^27,28^, cognitive tasks^29^, and motor tasks^30^. It is important to note that while encouraging, each of these studies suffers from one or more of the following: small sample sizes, selecting a region of interest (ROI) a priori, limited recording coverage only over those ROIs, correspondingly low channel counts, and a drop in reliability when the fNIRS cap/optodes were removed between measurements.

fNIRS comes in several flavors: continuous-wave (CW), frequency-domain (FD) and time-domain (TD). The latter (TD-fNIRS) is considered the gold standard of non-invasive optical brain imaging systems^31^. TD-fNIRS instruments can perform depth-resolved measurements, and provide improved and more quantitative estimates for both oxy- and deoxy-hemoglobin concentrations^32–34^. Until the development of Kernel Flow, this technology was not widely used and had been relatively inaccessible due to its bulky, expensive, and complex nature^34^. Kernel’s first-generation Flow headset was benchmarked using a set of standardized protocols for TD-fNIRS systems^35^ and validated in human studies^36,37^. Nonetheless, to our knowledge—barring a few reports^38^—very few studies to date have done a comprehensive TRR quantification of TD-fNIRS measurements and if so, they only focused on very limited features of the modality. To bridge this gap, the current study tested the validity and stability of Kernel’s second-generation portable and scalable neural recording system Flow2 in healthy subjects.

We hypothesized that Flow2 measurements could potentially go above and beyond the current state of fNIRS, TTR because: 1) the headset design enables reliable replacement after removal; 2) the headset boasts coverage over the whole scalp and provides high spatial, as well as temporal, resolution; and 3) recorded signals are more resilient to artifacts due the TD-fNIRS modality of the system. Leveraging the whole-head coverage of Flow2, we were able to not only demonstrate the consistency of brain activation patterns with prior literature, but also evaluate the stability of brain activity across different regions of the cortex. Moreover, with this approach, we confirmed that the observed TRR of neural activity was genuinely task-related rather than merely stemming from global signals, physiological factors, or artifacts. By employing a longitudinal multi-task experimental design, including a resting state, sensory, and cognitive task, we assessed the performance and stability of the Flow2 within an individual across time and across different headsets. Adding to the portable and scalable design of Flow2, this study provides a critical test of its clinical utility by quantifying its ability to reliably measure various neural activation features.

## Results

### Experimental design and data collection

The study was a repeated measures design with the primary objective of comparing brain activation patterns to the same tasks on two separate days. Participants were 49 healthy individuals (18 female; age=43.94 ±14.57, mean±STD) who completed two study visits (Methods).

The first study visit had the following stages: (1) participants performed a resting state session immediately followed by a passive auditory task; (2) the headset was taken off and participants rested during a portion of which they filled out surveys; (3) participants performed another resting state session and a Go/No-Go task. Neural recordings were done during stages (1) and (3) either using the same Kernel Flow2 device, or a different device (half of the participants were assigned to the same device, i.e. STAY, cohort). The second study visit followed the same structure, except the STAY cohort utilized the other Flow2 headset for this visit (Fig. 1a). This design was chosen to ensure we could investigate both the effect of repeated measurements and alternative headsets (Methods).

**Figure 1:**
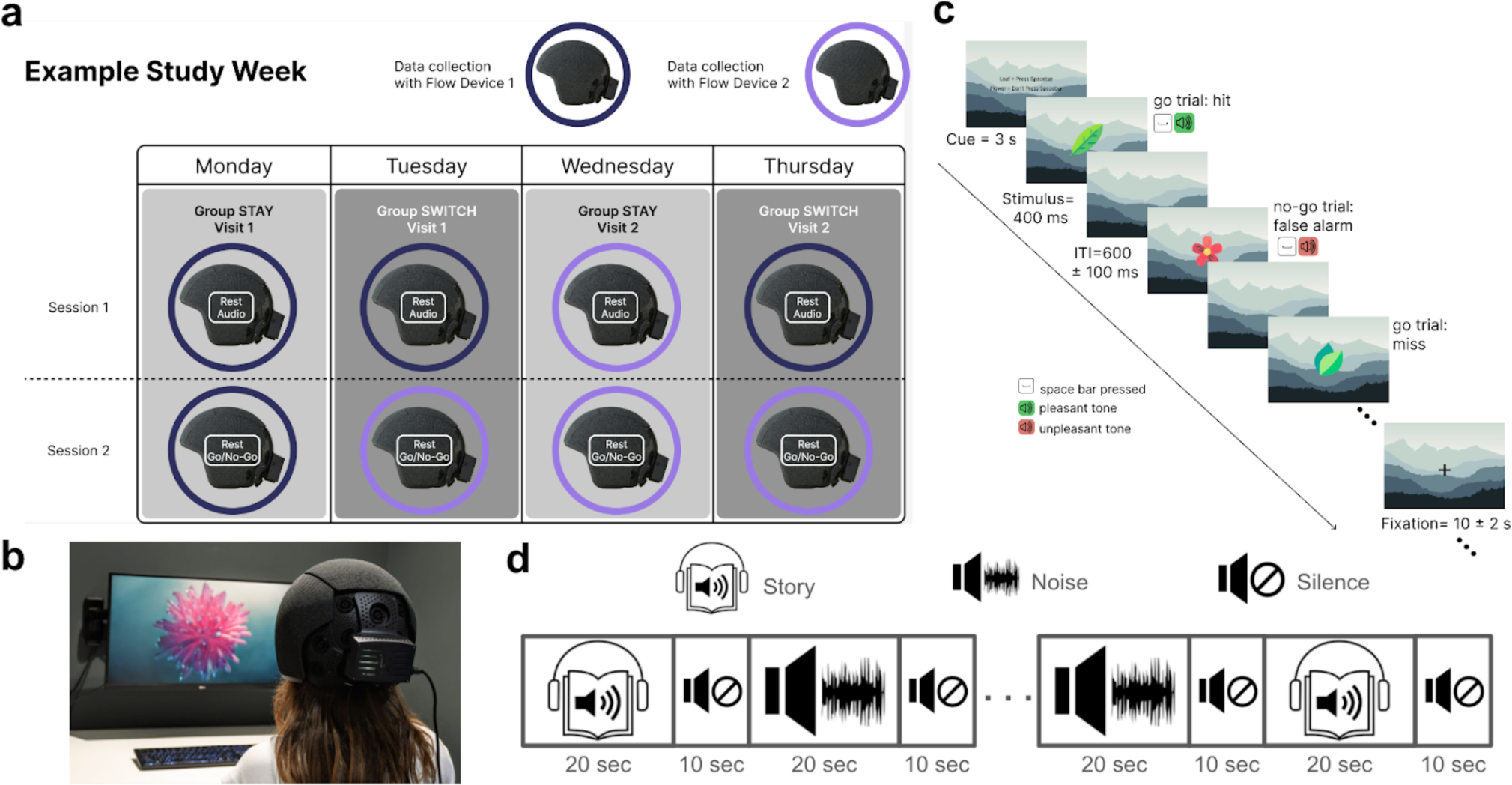
Overview of study design and experiments. **a.** Example schematic of a study week. Note that participants (either assigned to the group STAY or SWITCH) completed two study visits. Recordings during each visit were split into two parts: (1) resting state followed by a passive auditory task; and (2) resting state followed by a Go/No-go task. The headset was removed between these two stages. Different shades of purple denotes which headset was used for a given visit/session. **b.** A model wearing the Flow2 headset while performing the resting state session, which consisted of watching a 7-minute audiovisual segment. **c.** Schematic of the Go/No-Go task structure. Shown are a few representative trials at the start of a go/no-go block. **d.** Schematic of the passive auditory task with story and noise blocks (20s each) and 10s of silence in between each block.

During resting state sessions, the participants watched a 7-minute audiovisual segment^39^(Fig. 1b). The Go/No-Go task, which measures inhibitory control, consisted of blocks with low demand of control (go-only blocks) and high demand of control (go/no-go blocks)(Fig. 1c, Methods). Finally, the passive auditory task consisted of interleaved and pseudorandomized story blocks (20-second clips of TED talks) and noise blocks (20 second clips of brown noise)(Fig. 1d, Methods). During these tasks, we recorded hemodynamic signals using the Kernel Flow2 whole-head TD-fNIRS system (Fig. 2a).

**Figure 2:**
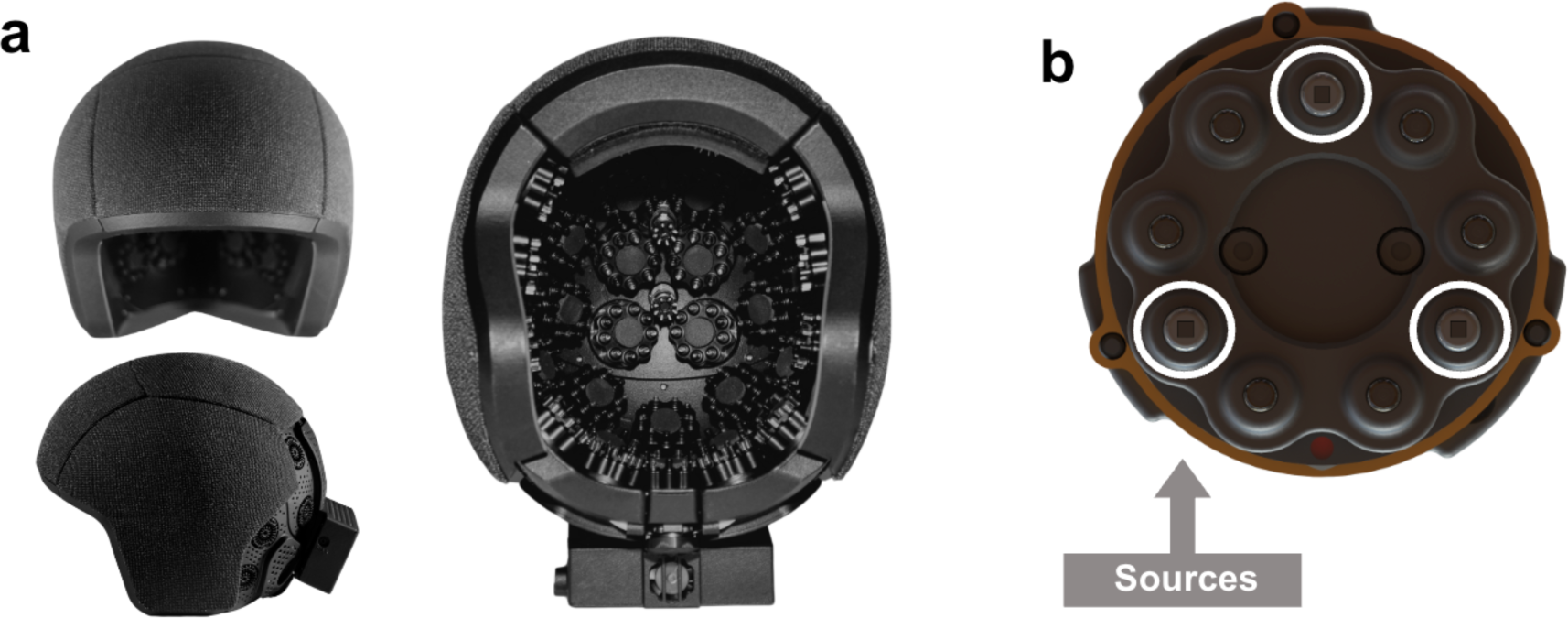
Kernel Flow2, second-generation Kernel whole-head TD-fNIRS system. **a.** Schematic of front, side and inside view of the Flow2 headset. Note the individual modules located throughout the headset thus providing whole-head coverage. **b.** Schematic of a module, which consists of 3 sources (marked by white circles) and 6 detectors.

The raw data from Kernel Flow2 consists of distributions of the time of flight of photons (DTOFs) collected for two wavelengths (690 and 905nm) from many source-detector pairs. Similar to Kernel Flow1, the Flow2 headset has a modular design, with a maximum of 40 modules (35 of which were utilized for this study), with each module containing 3 sources and 6 detectors (Fig. 2b). A channel is formed between a given source and a given detector. The different combination of within- and between-module channels can lead to thousands of channels with source-detector distances (SDS) between 8-60mm (Methods). Raw data from individual channels underwent standard preprocessing steps in order to obtain the first three moments of DTOFs (sum, mean, and variance), which were converted to the changes in the concentrations of oxygenated hemoglobin (HbO) and deoxygenated hemoglobin (HbR), as previously described^36,37^ (Methods). Moreover, we retrieved absolute chromophore concentrations with the curve fitting method, taking into account the instrument response function (IRF) which is continuously monitored for each source in the Flow2 headset^40^ (Methods). Several other features were subsequently computed, which will be described below (Methods).

In order to measure potential external sources of variability, participants were asked to fill out surveys (during stage (2)) detailing recent use of substances (e.g. caffeine, nicotine, etc) as well as daily activities (e.g., sleep satisfaction)(Methods). Only a small percentage of participants reported deviations in nicotine, alcohol, and marijuana use across the two visits (Supplementary Fig. 1a). A larger portion of participants reported changes in caffeine consumption and current sleepiness (change in caffeine consumption: 35%, change in sleepiness: 65%), though sleep satisfaction was still correlated between the two visits (⍴=0.43, p=2.25×10^−3^; Supplementary Fig. 1b). As such, these changes may affect brain activity, thereby altering across-day TRR metrics in the current study^41^.

#### Reliability of headset placement and coverage

A major source of variability in neurophysiological data in the current study could be due to the differences in headset placements during data collection and the resulting differences in optical channel locations. The inconsistency in placement can be both an issue within a given visit, as the headset was removed between stages (1) and (3) listed above, as well as between visits, i.e. across two different days. As such, AprilTags^42^ were placed on the headsets and participants were photographed each time the headset was donned (Supplementary Fig. 2a). Utilizing open-source computer vision algorithms^43^ and manual verification, we were able to obtain horizontal and vertical shifts (yaw and pitch dimensions) of the headset with respect to the participant’s head for each recording session (Methods; Supplementary Fig. 2b). This allowed us to compute the headset placement variability (i.e., standard deviation) for each participant across the 4 different placements. We showed that this variability is only on the order of a few millimeters (Supplementary Fig. 2c; horizontal shift: 3.23±1.70 mm, vertical shift:3.51±1.94 mm). This suggests that headset placement variability is lower than the spatial resolution of fNIRS (over 10 mm^44^), and may have minimal effect on data reliability, although a thorough quantification of this effect remains to be done.

Furthermore, we explored how reliably we could obtain similar levels of channel coverage in our recordings. After each headset placement, different sources and detectors could make good contact with the scalp, therefore leading to different numbers of analyzable channels (henceforth, referred to as retained channels; Methods), a potential confound in interpreting the reliability. Thus, we quantified this over multiple hierarchical scales (e.g., over the whole head, over the prefrontal region, and on an individual module level) as the fraction of retained channels in a given region (i.e. number of usable channels divided by the total number of possible channels). We then computed a Spearman correlation coefficient between within-visit and across-visit recordings. Fraction of retained channels, both in the prefrontal region (Supplementary Fig. 3a) and over the whole-head, was highly reliable in both within-visit and across-visit. This was indicated by a strong correlation value and proximity to the diagonal line (Prefrontal: within-visit ⍴=0.96, across-visit ⍴=0.94, p<10^−5^ for both; Whole-head: within-visit ⍴=0.99, across-visit ⍴=0.98, p<10^−5^ for both). Additionally, we had adequate coverage over the head as depicted by the number of retained channels from a given module (Supplementary Fig. 3b). This result suggests that variability due to different numbers of channels on different sessions may be minimal.

#### Reliability of resting state features across time and headsets

As noted above, participants completed a total of four resting state sessions (Fig. 1a). Our experimental design enabled us to consider both within-visit reliability (between two resting state sessions within a given visit) as well as across-visit reliability (between two resting state sessions across visits). Various features spanning physiological, optical and neural domains can be extracted from Flow2 headset recordings during resting state. In the current study, we focused on a non-exhaustive set of features, which were affected by pharmacological manipulation in our prior studies^37^, and/or which have strong literature support in terms of relating to individual differences. These features are from the four following categories: (1) absolute HbO/HbR in the prefrontal region, (2) effective attenuation coefficient (EAC)^45^ calculated at two different wavelengths (3) fractional amplitude of low frequency fluctuations (fALFF) within the left and and right prefrontal regions for HbO/HbR, (4) and functional connectivity (FC) within the left and right prefrontal regions for HbO/HbR (Methods).

Both within-visit and across-visit reliability for participant-level features were assessed using correlation analysis (Methods). First, we found that for both comparisons, many features of interest exhibited high reliability as evidenced by the strong relationship and proximity to the line of unity for these features (Fig. 3a). In fact, absolute HbO/HbR, EAC, and prefrontal fALFF showed strong correlation coefficients (>0.5) for both within-visit and across-visit reliability, while FC showed moderate to strong correlations (>=0.3), regardless of whether the same or different headset was used (data not shown). The results from all features for both within- and across-visit reliability are summarized in Fig. 3b. When considering reliability across all four resting state sessions, we computed other common metrics, namely the intraclass correlation coefficient (ICC) and Cronbach’s alpha. Both metrics demonstrated reliability above fair/acceptable for all features, and above good for most measures (Fig. 3c). Although FC within the prefrontal region showed the lowest reliability, we hypothesize that this may be due to the fact that this metric is more state-dependent (e.g. affected by mood, sleep, etc) compared to the other features.

**Figure 3:**
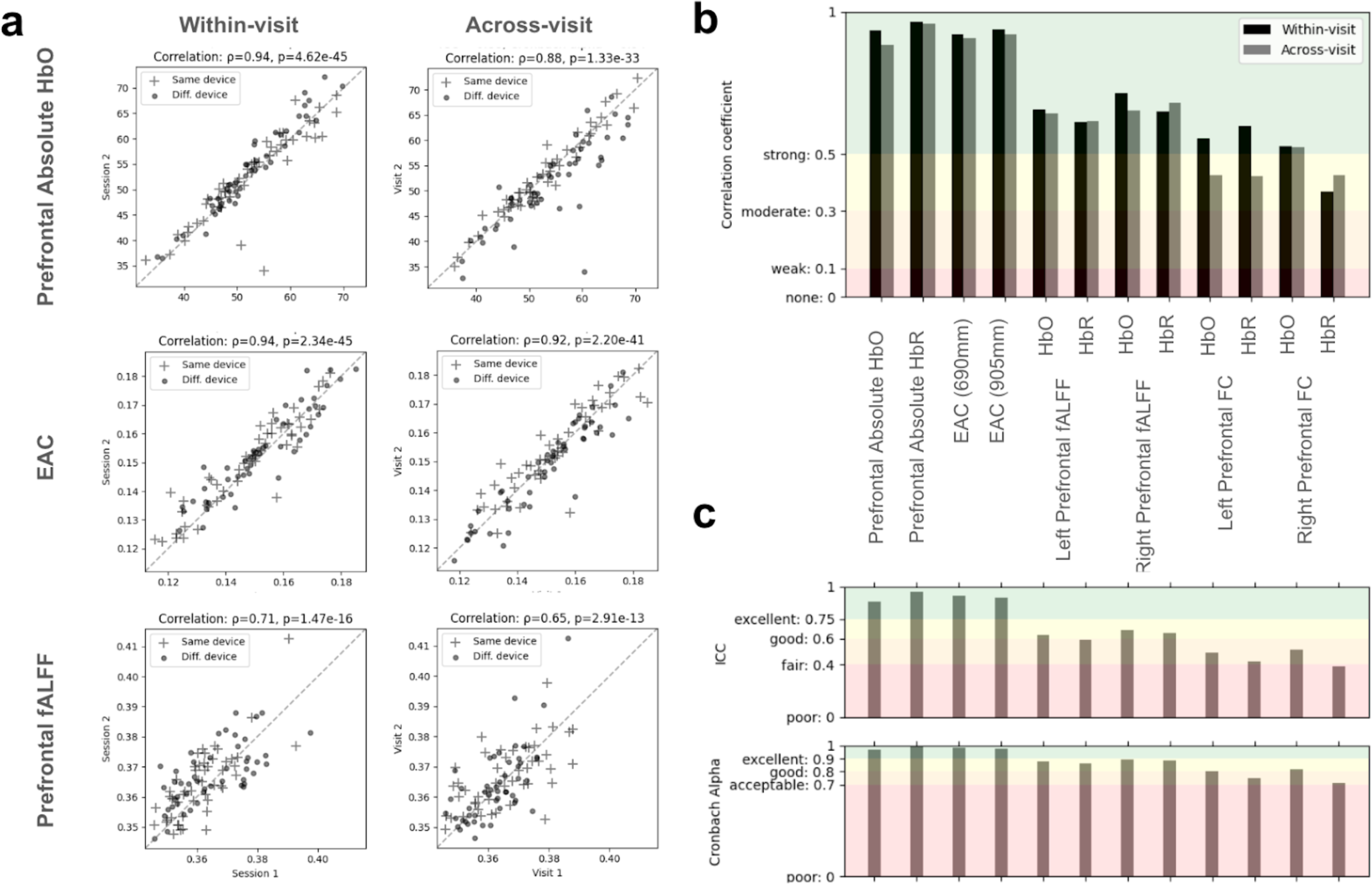
Various features revealed within-visit and across-visit reliability during resting state sessions. **a.** Different representative resting state features (each row) were highly correlated between two sessions within a given visit (i.e. between visit_1_session_1_ and visit_1_session_2_; and between between visit_2_session_1_ and visit_2_session_2;_ left column; within-visit) and across visits (i.e. between visit_1_session_1_ and visit_2_session_1_; and between visit_1_session_2_ and visit_2_session_2_; right column; across-visit). The three shown features are the absolute HbO [µM] in the prefrontal region, EAC of the 905 nm wavelength, and fALFF within the right prefrontal region HbO. Note that in addition to the high correlation coefficients, the values lie very close to the diagonal line (dashed line) indicating the similarity of the values. **b.** Within- and across-visit reliability (black and gray vertical bars respectively) as measured by the Spearman correlation coefficient between different visits/sessions, as described in (a), are shown for all resting state features. The colored background segments correspond to the commonly-used thresholds for the strength of correlation. **c.** Reliability of features across all four resting state sessions were computed using ICC (top) and Cronbach alpha (bottom). Colored segments depict different reliability thresholds as commonly used in the literature.

After investigating reliability in time, we wondered whether our recordings were reliable when using different headsets. We probed both within-visit and across-visit effects of the headset on the reliability of our features by comparing the difference in features between groups (i.e., same vs. different headsets; Methods). Of the twelve features of interest, and considering within-/across-visit effects, only the absolute prefrontal HbO in the across-visit showed potential headset related differences (independent t-test p<0.05; corrected). Despite this difference, features from both same and different headset lie very close to the line of unity, and exhibit high reliability as measured by the correlation coefficient (HbO across-visit: same headset ⍴=0.96, different headset ⍴=0.83). Additionally, it is unlikely that this observation is purely driven by the use of different headsets as the within-visit analysis revealed no significant differences. Taken together, the explored resting state features were adequately reliable despite factors such as the passage of time and using different headsets.

While we could not investigate all possible confounding variables that could affect reliability—in addition to considering the effects of headset placement, channel coupling, time (within-visit/across-visit), and different headsets—we also considered participants’ skin color (Methods). First, we found that skin color had a significant effect on the total number of retained channels in the prefrontal area (Supplementary Fig. 4a). Based on the line of best fit, it is anticipated that prefrontal channel count at the highest melanin levels will still exceed 200, providing ample coverage for computing the neural features reported in this study. Reinforcing the assertion of the practical insignificance of this relationship, prefrontal absolute HbO was not correlated with skin color (Supplementary Fig. 4b). This is particularly important after the COVID-19 pandemic revealed systematic deviations in blood oxygenation levels as a function of skin color^46^. Finally, we performed a median split on participants’ forehead melanin levels and probed the reliability of absolute oxygenation for low and high melanin groups separately. We found high reliability across visits for both groups (low melanin: Pearson r=0.86, p=5.79×10^−8^; high melanin: Pearson r=0.94, p=2.55×10^−12^; Supplementary Fig. 4c). Absolute deoxygenation also exhibited similar levels of high reliability for both groups (low melanin: Pearson r=0.93, p=5.93×10^−11^; high melanin Pearson r=0.96, p=5.94×10^−14^).

#### Reliability of hemodynamic activity during sensory (passive auditory) task

Participants completed a passive auditory task twice (once in each study visit), where half of the participants donned the same Flow2 device, while the other half switched devices (Fig. 1a, d). To investigate brain activation patterns associated with this task, we employed a Generalized Linear Model (GLM) framework, where the activity of each channel was modeled as a function of block type (story and noise) and other nuisance regressors^36^. To obtain group-level activations, for each channel we pooled the fitted model weights (beta values) associated with the block conditions and performed a one-sample t-test (Methods). We repeated this process for visit 1 and visit 2 separately. Importantly, we found that the expected regions (auditory cortices) were activated during the story condition of the task^47,48^. This activation was apparent in the group-level GLMs for both visit 1 and visit 2, as the foci of significant activations were in the lateral temporal regions (Fig. 4a).

**Figure 4.**
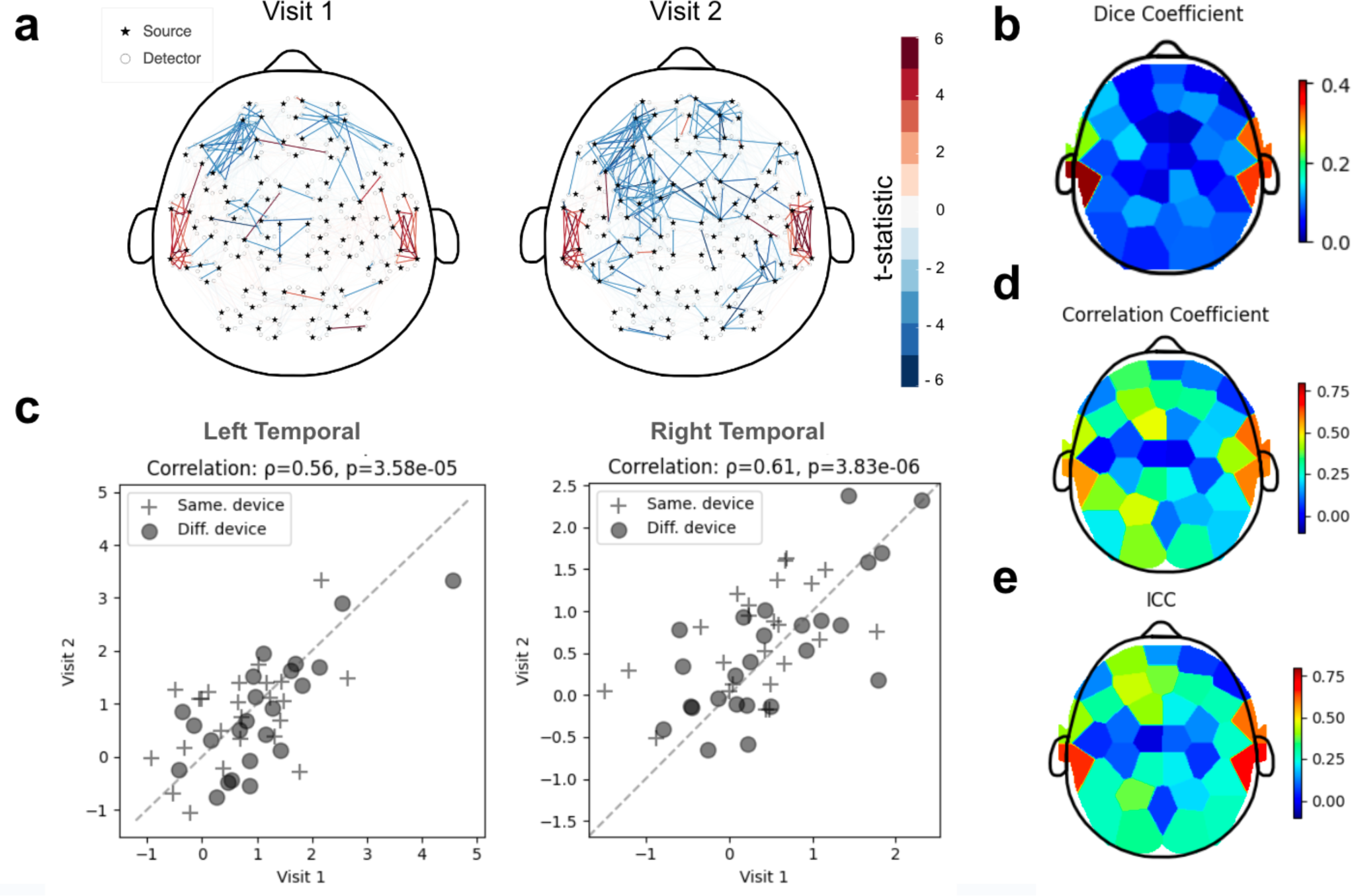
Activations in the auditory cortex during the passive auditory task were reliable across study visits. **a.** Group-level GLMs for the story condition during visit 1 (left) and visit 2 (right) were qualitatively similar as evidenced by the t-statistics from a one-sample t-test. Here, for each channel (each line) GLM beta values from all participants were compared against zero. Only channels that were significantly different from zero (p<0.01; uncorrected) are shown. **b.** The group-level dice coefficient (a measure of channel-level reliability) was computed as the average dice coefficient of all participants for each module (each color patch). Note the patches with high values (>0.4) in the auditory cortex areas. **c.** For each participant (each marker), the average GLM test statistics for each module was calculated and compared between two recordings (i.e. visit 2 GLM test statistics versus visit 1 GLM test statistics). Two representative modules in bilateral auditory regions showed strong and significant correlations (Spearman) between the two visits. **d, e**. Correlation coefficients (Spearman ⍴) (**d**) and ICC values (**e**) computed to measure stability of GLM test statistics (across time) are shown for each module (each color patch). Note how the reliability as measured by dice coefficient (**b**), correlation coefficient (**d**) and ICC (**e**) exhibit consistent patterns, with the left and right auditory areas showing the highest reliability (bluer patches indicate lower reliability and redder patches indicate higher reliability). Results shown here are all computed over the story condition of the task.

First, we quantified the reliability of these activations at the resolution of individual channels by computing the dice coefficient, a quantitative measure of the similarity (or overlap) between two sets (Methods). Applied here, the two sets of interest were the significantly activated channels detected by a given module in Visit 1 and those in Visit 2. For reference, sets containing the exact same elements yield a dice coefficient of 1, and sets with no common elements yield a dice coefficient of 0. We computed the dice coefficient for each participant and each module, obtaining whole-head dice coefficient maps. The averaged (across participant) dice coefficient map showed elevated measures in the auditory cortex similar to those reported in prior fNIRS literature^27^, whereas all other regions exhibited averages near-zero (Fig. 4b).

Next, in order to assess the reliability of the magnitude of task activations, we employed a module-level approach and averaged participant-level GLM test statistics over all channels detected within a given module. For each module, we computed the correlation between the participant-averaged test statistics from visit 1 and that from visit 2 (Methods). This analysis revealed that the magnitude of participant-level activations in bilateral auditory cortex modules were highly correlated across the two visits (Left Temporal ROI: ⍴=0.56, p=3.58×10^−5^; Right Temporal ROI: ⍴=0.61, p=3.83×10^−6^). Importantly this relationship was close to the line of unity (Fig. 4c) indicating consistent activation (i.e. no general increase or decrease in activation between visits). Similar to the resting state features, the change of headset did not have a significant effect on this reliability (independent t-test; p>0.05 for all modules). In addition to Spearman correlation, we also computed the ICC on these same module-averaged values and visualized them over the whole head (Fig. 4d, e). In accordance with the dice coefficient analysis, we found auditory areas to have the highest reliability both in terms of correlation coefficients and ICC values (Left Temporal ROI: ICC=0.65; Right Temporal ROI: ICC=0.70). In fact, both values exceed the recommended interpretation thresholds indicative of strong relationships (r > 0.5) and good agreement between measurements (ICC > 0.6). With these findings in mind, Flow2 can measure reliable brain activations resulting from sensory stimulation.

#### Reliability of hemodynamic activity during cognitive (inhibitory control) task

Participants were asked to perform a commonly employed inhibitory control task (Go/No-Go) during each study visit. As with the auditory task, we utilized different Flow2 devices for half of the participants (Fig. 1a, c). By definition, cognitive tasks are more complex than sensory tasks, and reliability can be affected by behavioral complexities such as strategic and attentional shifts. Nevertheless, we sought to measure the reliability of brain activity while participants performed the Go/No-Go task. In an attempt to guard against major changes in performance (and thus corresponding changes in brain activity), participants were given ample time to practice the task prior to the first recording, and behavioral data (accuracy and median reaction time in go/no-go blocks) was also collected (Methods).

Indeed, behavioral metrics were correlated across the two visits, such that high performers from visit 1 tended to be high performers in visit 2. These relationships (accuracy: ⍴=0.77, p=9.81×10^−11^; reaction time: ⍴=0.60, p=6.52×10^−6^) were fairly strong (Supplementary Fig. 5) indicating that behavior was stable enough to expect a reflection of that stability in the neural activity. While strong, it is important to note that any change in performance should lead to ‘less stable’ brain measurements, as we expect task-related Regions of Interest (ROIs) to be sensitive and responsive to behavioral fluctuations. In this way, TRR of task performance, sets an upper bound for TRR of neural metrics.

For this task, we similarly fitted a GLM to the neural activity from the recordings and performed the same analyses (here, the block types were go-only and go/no-go). Group-level analysis on the GLMs for visit 1 and visit 2 revealed significant activations in right prefrontal regions during the go/no-go blocks, a result that is in agreement with this area’s role in inhibitory control^36,49–51^. With whole-head coverage we observed significant activations in other areas, such as the right auditory cortex and the left auditory/motor regions (Fig. 5a.). The noted auditory activation is consistent with the trial-level auditory feedback (Methods). One possible explanation for the left motor activation is the propensity of right-handedness in participants (88%), who were asked to respond with spacebar presses in this task.

**Figure 5.**
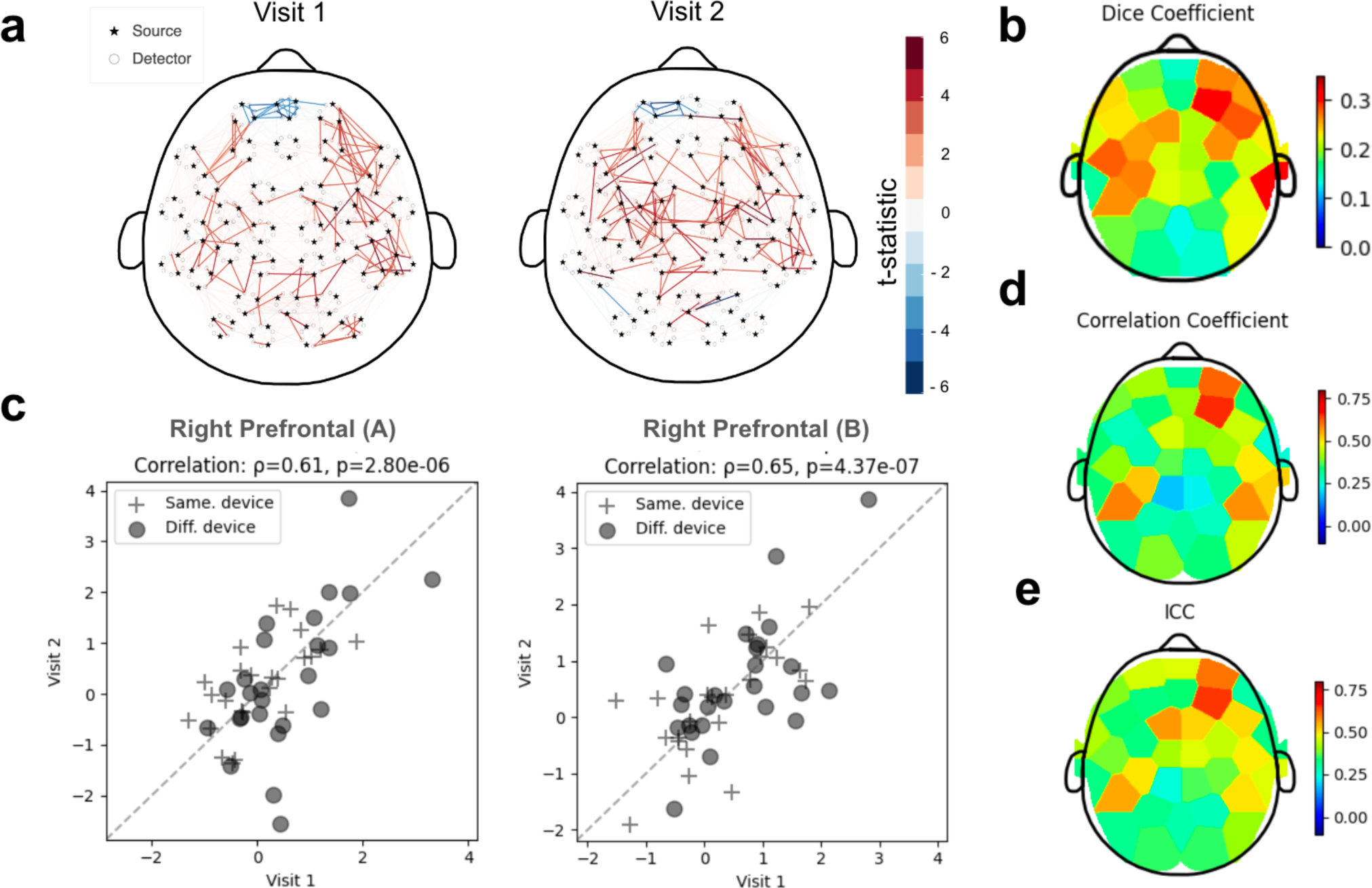
Reliability of the right prefrontal regions was observed in a Go/No-Go inhibitory control task. **a.** Shown are the t-statistics from a one-sample t-test on channel-wise GLM beta values for each participant, for the contrast go/no-go. These group-level GLMs showed similarity in brain activation patterns during the task between visit 1 (left) and visit 2 (right). Only channels that were significantly different from zero (p<0.01; uncorrected) are shown. **b.** The dice coefficient of all participants were averaged for each module to obtain the group-level dice coefficient (a measure of channel-level reliability). Each patch represents a module. Note the presence of patches with high values in the following areas: right prefrontal, right auditory, and left auditory/motor. **c.** Two representative modules in the right prefrontal region showed strong and significant correlations (Spearman) between the two visits. Each marker represents the average GLM test statistics for each module for a given participant compared between two recordings (i.e. visit 2 GLM test statistics versus visit 1 GLM test statistics). **d, e**. Module-level reliability of GLM test statistics (across time) is shown using correlation coefficient (Spearman ⍴) (**d**) and ICC values (**e**). Each color patch represents a module. Several regions, and primarily the right prefrontal region, showed high reliability consistently across all three measures of reliability (**b, d, e**). Results shown here are all computed over the go/no-go condition of the task. In all heatmaps bluer patches indicate lower reliability and redder patches indicate higher reliability.

Subsequent dice coefficient analysis revealed elevated reliability in each of these regions at the population-level, indicating consistent channel-level activations in these regions across the two visits (on average) (Fig. 5b). Correlation analysis on the GLM test statistics revealed a strong agreement within ROIs in the right prefrontal cortex (Right Prefrontal Module A: ⍴=0.61, p=2.80×10^−6^; Right Prefrontal Module B: ⍴=0.65, p=4.37×10^−7^)(Fig. 5c, d). In accordance with the auditory task, none of the ROIs showed significant differences in reliability when using the same or different headsets for recordings (independent t-test; p>0.05 corrected). Last, ICC was computed over the module-level GLM test statistics (Fig. 5e) and the same two modules positioned over the right prefrontal cortex showed the highest ICC values (Right Prefrontal Module A: ICC=0.60; Right Prefrontal Module B: ICC=0.64)(Fig. 5e). While evoking and measuring reliable brain activity as it pertains to cognitive tasks has its challenges, here we showed that brain activity recorded by Flow2 produced reliable brain metrics that were on par with the stability of the task performance across visits and consistent with prior fMRI literature^52^.

## Discussion

The present study examined the reliability of neuroimaging measurements in participants performing resting state, sensory and cognitive tasks while their data were recorded using Kernel Flow2 TD-fNIRS system. To achieve this, we incorporated multiple factors that could lead to measurement variability into our experimental design, such as (1) passage of time between measurements; (2) using a different system for data acquisition; and (3) variability in headset placement and its consequences (e.g. different number of useable channels). We demonstrate valid and stable measurements across several neural features computed from tasks and resting state Flow2 recordings.

By extracting a combination of neural, physiological and optical features from the resting state sessions, our data revealed a reliability in the good to excellent range for most investigated features across standard reliability metrics (e.g., correlation coefficient, ICC and cronbach alpha). This reliability was observed both across time and the use of different headsets. It is worth noting that although our reliability metrics were comparable, and at times superior, to those reported in fMRI^10,11,13^ and fNIRS^25^, other studies may have used different metrics to evaluate reliability. Additionally, the limited number of features used to assess the reliability of resting state in the current study were spatially crude (e.g., fALFF over the entire right prefrontal region). It is possible that reliability metrics would improve, or deteriorate, if more granular ROIs are considered—an investigation worth pursuing in future studies.

Reliability of neural activations during tasks has been a topic of interest in both fMRI^12^ and fNIRS^27–30^ communities. Although one expects to observe very stable neural responses during sensory tasks^27^, which was indeed the case in our passive auditory task, activation stability during cognitive tasks is more nuanced. Even in the absence of any neural recordings, behavioral performance during cognitive tasks and cognitive scores can vary from day to day^52,53^. In parallel, it is well-known that measured brain activations in the prefrontal regions can be related to performance on Go/No-Go tasks^36,49–51^; As such, it is possible that the variability in behavioral performance, albeit inevitable, may impose an upper bound on the stability of corresponding neural responses during cognitive tasks. Even still, in the current study we found adequately high reliability during our Go/No-Go task that matched inhibitory control reliability in fMRI^52^; and provided a more thorough analysis by utilizing complementary metrics.

When assessing task-based reliability, we examined a wide range of metrics to ensure consistency across these metrics. For example, in conjunction with the traditional metrics such as correlation and ICC, we also considered the dice coefficient. Even though we obtained confirmatory results between all our metrics, we believe they can paint different pictures of how reliability is measured. While the dice coefficient can provide a great metric for spatial stability of responses, the correlation and ICC values provide a complementary metric that also incorporates the magnitude of responses in each region. Furthermore, most fNIRS-based TRR studies only consider *a priori* defined regions of interest that are thought to be implicated in the task^27–30^ due to the limited number of available optodes. One of the advantages of the whole-head Flow2 system is that it allows for probing reliability beyond these predefined regions. Here, we were able to determine that brain activation patterns and stability varied across the cortex with task-related regions showing the strongest measures – an important validation that system measurements are being driven by brain activity (and not noise or global signals). Outside of the expected region of interest, we observed an area in the auditory region that exhibited high reliability in the Go/No-go task, which we believe may reflect a response to the auditory feedback participants received during the task.

It must be borne in mind that the current study, similar to other TD-fNIRS studies, has multiple limitations. First due to the nature of this modality, it is only possible to report reliability in superficial brain regions. Second, our study design measured test-retest reliability over one week, so we cannot report on reliability over longer timescales, and we did not explore systematic changes in underlying metrics due to factors like circadian rhythms or seasonal changes^54^. Third, our post-hoc analysis revealed a statistically significant relationship between melanin and total number of retained prefrontal channels; however, the practical significance of this relationship was demonstrated to be negligible as anticipated channel count (> 200 across skin colors) was more than adequate for the extraction of the neural metrics considered here (e.g. the absolute HbO/HbR metric). Still, our current study was not specifically designed to address the effect of skin color on our recordings and we did not make recruitment decisions based on this factor. As such the participants were not balanced across melanin levels, which could affect the interpretability of the analyses we performed; nonetheless, our recruited demographics were similar to that of the Greater Los Angeles area (data not shown). Lastly, we know that hair color and texture can also affect fNIRS recording quality. Although, on average, we had adequate number of channels over the head, a proper quantification of how this relates to hair texture remains to be done in future studies.

To summarize, quantifying the test-retest reliability of the metrics derived from a neuroimaging system is critical. If a system were to be used for diagnostic purposes and identification of neuropsychiatric and/or neurodegenerative biomarkers, a lack of adequate reliability could obscure any true underlying signals^1^. The reliability of signals recorded from Kernel Flow2—on a variety of tasks, at multiple time points, and using different headsets—suggests its potential to fill a need for scalable multi-device and even multi-site neuroimaging in the scientific and medical communities.

## Methods

### Participants and screening procedures

Forty nine healthy participants (18 female, age=43.94±14.57, mean±STD) completed two study visits in a repeated-measures study design within a week (time between visits: mean±STD=2.02±0.78 days, range=1–6 days)(Fig. 1a). Inclusion criteria were: (1) healthy adults who are 18 years or older at the time of enrollment; (2) must have the ability to consent for themselves; (3) must be fluent in English (speaking and reading), and (4) must be willing to attend planned study visits at the research site. Exclusion criteria were: (1) major visual or auditory deficits that would prevent them from completing a study task; (2) being or the possibility of being pregnant (for people of childbearing potential [POCBP]); (3) any history of severe neurological or severe psychiatric disorders, including head trauma with serious results (coma, unconscious for >2 hrs, or skull fracture); (4) any other psychiatric disorder with unstable treatment in the prior 6 months; (5) lifetime substance or alcohol dependence, and/or alcohol or substance abuse as determined by CAGE-AID assessment for drug and alcohol abuse^55^ (a score of 2 or higher would result in exclusion); (6) current or recent (in the past 6 months) chemotherapy and/or radiation for any cancer; (7) hospitalizations and/or unstable health/medical condition/treatment in the last 30 days; and (8) does not agree to image recording, including photographs, videos, and/or 3D scans of the head and face, during the study.

Participants gave written informed consent before beginning the study in accordance with the ethical review of the Advarra IRB (#Pro00074416), which approved this study, and the Declaration of Helsinki. All experimental protocols were approved by the IRB. Participants received monetary compensation for their time, effort, and travel expenses.

In this study 49 participants are included in the analysis. We recorded data from an additional 6 participants, but excluded them from analysis for the following reasons: missed visit (n=2), headset discomfort (n=2), gum chewed during recording (n=1), and desire to not remove head bandana for recording (n=1). All Flow2 data was rigorously checked for quality before being included in data analysis. Importantly, no data was lost due to technical issues; therefore all 49 participants are included in all of the analyses. For the cognitive task, performance was adequate to ensure that participants understood the task in all cases.

### Study design

#### General description

The study was a repeated-measures randomized design with participants completing two study visits within a week (Fig. 1). To measure test-retest reliability (TTR), participants completed the following sequence during visit 1: (i) a resting state session; (ii) a passive auditory task; (iii) filling out surveys + misc (see below); the headset was removed during this phase; (iv) another resting state session; and (v) a Go/No-Go task. Participants’ brain activity during stages i, ii, iv and v was recorded with Kernel Flow2 Time Domain functional Near-infrared Spectroscopy (TD-fNIRS). The second study visit followed the exact same structure.

During their first visit, a few additional steps were completed. Prior to data collection, participants listened to all audio clips included in the passive auditory task to eliminate the effect of novelty on neural signals during the experiment. Furthermore, during phase (iii), participants were trained on the Go/No-Go task by practicing on six shortened blocks (15 trials instead of the original 24 trials), three with go-only trials and three with both go and no-go trials, in attempts to remove learning effects from the neural recordings. During phase (iii), we recorded the melanin index^56^ on the parietal regions of the scalp/hair as well as on the forehead using a colorimeter (Cortex Technology DSM III Color Meter, CyberDerm, Broomall, PA). Only the data from the forehead readings, which were considered to be pure measures of the skin color (unoccluded by any hair), were used for further analyses. Lastly, participants were asked to fill out a self-report demographics questionnaire part of which includes the Fitzpatrick scale for skin typing.

In addition to investigating reliability over time, we sought to explore the reliability across two different Flow2 headsets. To this end, participants were randomly assigned to one of the following two groups (Fig. 1a):

1. STAY group: during the first visit, headset 1 was used for both the first half (i,ii) and second half of the visit (iv, v); during the second visit, a different headset, i.e. headset 2, was used for both halves.
2. SWITCH group: during the first visit, headset 1 was used for the first half (i,ii) and headset 2 was used for the second half of the visit (iv, v); during the second visit, the same headset order was used.

This study design allowed for investigation of within-visit reliability with the same headset, within-visit reliability with different headsets, across-visit reliability with the same headset, and across-visit reliability with different headsets.

At each visit, participants filled out a survey during phase (iii) that asked about recent daily activities, including: quality of sleep (scale 1–10), current sleepiness level, caffeine consumption (within the last 12 hours), nicotine use (within the last 3 hours), alcohol consumption (within the last 12 hours), and marijuana (within the last 72 hours). With the exception of sleep quality all of the questions were qualitative multiple choice.

#### Go/No-Go task paradigm

The task, designed and presented using Unity game engine, consisted of two block types: go-only and go/no-go. The overall structure of the task was similar to the one used in a prior publication^36^. Briefly, participants completed a total of 10 blocks alternating between go-only (n=5) and go/no-go (n=5) with each block consisting of 24 trials. Stimuli were green leaf cartoon images (for go trials) and red flower images (for no-go trials) that were presented in a pseudorandom order, which was pre-set and unique for each run. During go/no-go blocks, 30% of the trials were chosen to be no-go trials. A different run of the task was presented at each study visit, however, all participants completed the same versions in the same order.

The task included a 15-second rest at the beginning and a 20s rest at the end. The task also included a screen to remind the participant of task instructions (e.g. pressing the spacebar when seeing a green leaf and refraining from pressing when seeing a red flower) prior to each block. Stimulus presentation time was 400 ms followed by a 600ms±100ms inter-trial interval during which only the background was displayed. Participants were asked to provide a response within the 400ms presentation period. A pleasant or unpleasant tone, played immediately after participants’ response, was used to provide positive (for hits) and negative feedback (for false-alarms), respectively. The overall task duration was approximately 7 minutes (Fig. 1c).

#### Passive Auditory task paradigm

This task was also programmed and presented in Unity. The design consisted of two types of blocks, each lasting for 20s: story blocks (n=8; 6 unique clips and 2 repeated clips) during which participants listened to short clips from TED talks; and noise blocks (n=7) during which participants listened to brown noise. After an initial 10s rest period, the story and noise blocks were presented in a preset pseudo-randomized order with 10s of silence between each block (Fig. 1d). Participants wore earbuds and were instructed to look at a white fixation cross presented on a black background throughout the task. This task required no cognitive demand and was purely a passive auditory task. All participants listened to the same version of the task (i.e. in terms of the audio clips and their order) in both visits, with the exception of three participants who listened to a different version of the task in the second visit.

### Headset placement analysis

This procedure involved multiple steps: (1) four distinct AprilTags^42^ were installed on each headset in fixed locations (Supplementary Fig. 2a); (2) A front-facing photograph of each participant was taken each time the headset was placed on their head (immediately before data collection started), yielding four pictures per participant; (3) Finally, a computer vision (CV) approach^43^ was used to calculate horizontal and vertical shifts of the headset relative to facial features, and in turn within-participant variability of operator headset placement.

For processing, each photo was converted to a JPG, resized to a standard pixel dimension, and converted to grayscale. Two python packages were used to implement facial feature detection and AprilTag detection: The dlib library was used to detect faces and facial features in images (68-landmark shape predictor model), and the Apriltag library was used to detect AprilTags in the images. Detection results included tag corners and center coordinates. These locations were then used to quantify alignment of the headset as follows (Supplementary Fig. 2b): i) horizontal shift (yaw) was measured as the horizontal distance between the nose-line (defined by the landmarks detected along the nose) and the center of the two frontal, innermost april tags, and ii) vertical shift (pitch) was computed as the average length of the line connecting the center of each eye to the AprilTag above it. All metrics are reported in millimeters.

Photographs were missing from 4 of the 196 placements. Of the remaining 192 photos, 140 photographs across 42 participants were included in the analysis because a subset of photos could not be analyzed. Some of these photos were excluded due to failure to detect necessary AprilTags (n=44), or due to camera angle (n=1). After the previous exclusions, 7 participants had only one analyzable photograph, which was insufficient for reliability calculations; therefore, their photos (n=7) were also removed from the analysis. Manual verification of facial landmark locations was performed on 44 of the analyzed photos, due to failure of the CV algorithm to accurately detect facial landmarks. The photos were taken head-on with a fixed position front-facing iPhone camera. Therefore, for ease of computation we assumed a simplified model that neglects the effects of curvature. While this assumption might introduce slight deviations in our results, these are anticipated to be minimal and do not significantly impact the overall conclusions within the context of our study. Results are shown in Supplementary Figure 2c.

### fNIRS data collection and feature extraction

#### Data acquisition

Participants wore the Kernel Flow2 TD-fNIRS headset (the second generation of Flow, Kernel, Culver City, CA, USA)^35,37^ for neurophysiological recordings. The system collects distributions of the times of flight of photons (DTOFs) from more than 3000 channels across the head with source-detector distances spanning 8-60mm. The system uses two wavelengths (690 nm and 905 nm) and samples at an effective rate of 3.76 Hz. The modular design of the headset can allow up to 40 modules arranged to provide coverage over prefrontal, parietal, temporal, and occipital regions across both hemispheres (Fig. 2a). Each module consists of three sources and six detectors yielding both within-module channels (up to 18)(Fig. 2b) as well as between module channels (variable number depending upon module location). For this study, we used a 35-module system by removing the most posterior modules that covered inferior occipital regions, which were deemed to be the least relevant to our tasks.

#### Data preprocessing and relative hemoglobin concentrations

Data preprocessing steps are described at length in prior work^37^. We first implemented a channel selection procedure using histogram shape criteria^35^. Next, histograms from the selected channels were used to compute the moments of the DTOFs–specifically the sum, mean, and variance moments^57,58^. The relative changes in preprocessed DTOF moments were converted to relative changes in absorption coefficients for each wavelength using the sensitivities of the different moments to changes in absorption coefficients (moments Jacobians, i.e. derivative of each moment with respect to a change in absorption coefficient), following the procedure recently described in ^33^. To obtain the sensitivities of the different moments, we used a 2-layer medium with a 12-mm superficial layer. We used a finite element modeling (FEM) forward model from NIRFAST^59,60^, and integrated the moment Jacobians within each layer to compute sensitivities. We converted the changes in absorption coefficients at each wavelength to changes in oxyhemoglobin and deoxyhemoglobin concentrations (HbO and HbR, respectively) using the extinction coefficients for the two wavelengths and the modified Beer–Lambert law (mBLL)^61^. HbO/HbR underwent further preprocessing using a motion correction algorithm, Temporal Derivative Distribution Repair (TDDR)^62^. TDDR can leave spiking artifacts in the presence of baseline shifts; we detected those and corrected them using cubic spline interpolation^63^. Finally, we detrended the data using a moving average with a 100-second kernel and applied short channel regression to remove superficial physiological signals from brain activity^64^ (here, short within-module channels with SDS=8mm were used).

#### Absolute hemoglobin concentrations

The DTOF is the convolution between the time-resolved temporal point spread function (TPSF) and the Instrument Response Function (IRF). We leveraged Flow2’s online IRF measurements (dedicated detector within a module, which records photons that come directly from the laser), and used a curve fitting procedure to retrieve absolute optical properties of the underlying tissue. We used an analytical solution of the diffusion equation for a semi-infinite homogeneous medium to generate candidate TPSFs, which we convolved with the known IRF, and compared with the recorded DTOF; the search for optical properties was conducted using the Levenberg-Marquardt algorithm, focusing on the fit in the region that spans from 80% of the peak on the rising edge to 0.1% of the peak on the falling edge. The refractive index was set to 1.4. This provided an approximate estimate of absorption coefficients as the semi-infinite assumption is an idealization. These absolute estimates of absorption coefficients were converted to HbO and HbR concentrations, as described above. To obtain a single value for HbO and HbR, we computed the median value across well-coupled long, within-module prefrontal channels (SDS=26.5mm) in the first 30s of a given resting state session.

#### Extraction of optical properties

During resting state sessions, we sought to measure head tissue opacity to near-infrared light, which provides information about the health status of the brain’s cortical mantle. We computed the Effective Attenuation Coefficient (EAC)^45^ using the slope of the log(SNR) of the signal, here the total counts (i.e., the sum moment) of DTOFs, as a function of source–detector distance.

#### Local neural activity: fractional Amplitude of Low Frequency Fluctuations (fALFF)

The time series of data from each channel (for each chromophore) during resting state sessions was converted to the frequency domain using an FFT. The ratio of the power in the 0.01–0.08Hz frequency range was calculated relative to the full 0–0.25Hz frequency range. We focused our analyses on left and right prefrontal regions only. As such, fALFF values for each chromophore were computed by averaging all left/right prefrontal channels.

#### Functional Connectivity (FC)

To obtain FC during resting state sessions, the time series of each channel was further processed by applying a low-pass filter (an acausal finite impulse response (FIR) filter). The pass band was 0.01-0.1Hz, as is customary in the literature^65^. Functional connectivity was computed for all pairs of channels as the Pearson correlation coefficient between their time courses, for each chromophore independently. Similar to fALFF, we focused on left/right prefrontal regions only. Values from the FC matrix within each of these regions were then averaged for each chromophore.

#### Generalized Linear Model (GLM) approach

The approach we employed was similar to methods previously described^36^. Each of the measured time courses, y (here, channel-wise relative HbO), were modeled as a function of a design matrix, X (a set of task specific regressors), and residuals, ϵ, as follows: y = Xβ + ϵ, where the β coefficients represent the contribution of each regressor to the data. This multiple linear regression problem is then solved using a least-squares solution that includes prewhitening the time courses with an autoregressive model. The design matrix, X, included a combination of block-level and standard nuisance regressors such as (1) time course of the task blocks (square waves convolved with a a canonical hemodynamic response function for each block type); (2) drift regressors; and (3) low frequency cosine terms. GLM Contrasts (test statistics) and their corresponding significance (p-value) were then computed between conditions of interest via statistical tests (t-tests) on the fitted β coefficients. We used this approach for both the passive auditory and Go/No-Go tasks.

1. *<u>Passive auditory task</u>*: Given that there were two types of blocks in the experimental design, story and noise blocks, the four possible contrasts were: story, noise, story - noise, and noise - story. For the purpose of this manuscript, we only considered the “story” contrast.
2. *<u>Go/No-Go</u>*: Similar to the audio task, there were two block types in the experimental design, go-only and go/no-go blocks. As such, the four possible contrasts were: go-only, go/no-go, go-only - go/no-go and go/no-go - go-only. For the purpose of this manuscript, we only considered the “go/no-go” contrast.

### Statistical analyses

#### Resting state

Note that for each participant, there were a total of four resting state sessions. Two were done during the first visit (referred to as sessions 1a and 1b) and the remaining two during the second visit (referred to as sessions 2a and 2b). For each session, we considered the following features: (1) Absolute HbO/HbR in the prefrontal region; (2) EAC for each wavelength; (3) fALFF computed for HbO and HbR in left and right prefrontal regions separately; and (4) FC matrix computed for HbO and HbR averaged within left and right prefrontal regions separately. Additionally, two different headsets were used for resting state recordings; for a detailed description, see Study Design section as well as Fig. 1a.

1. *<u>Pairwise Correlations</u>*: We considered both within-visit relationships, i.e. between sessions 1a/1b and between sessions 2a/2b as well as across-visit relationships, i.e. between sessions 1a/2a and sessions 1b/2b. For each of these pairs, a Spearman correlation coefficient was computed for each feature of interest. The results are shown in Fig. 3a, b.
2. *<u>Intra-class correlations and Cronbach’s alpha:</u>* In addition to pairwise correlations, we used the features from all four resting state sessions and computed the Intra-class correlation coefficient (ICC) as well as Cronbach’s alpha, both of which are commonly used metrics for reliability. We used the pingouin Python package^66^ for both ICC (specifically ICC2, known as the single random rater method) and Cronbach’s alpha calculations (participant IDs were used as targets/subjects variables and session IDs were used as raters/items variables for ICC/Cronbach alpha). Participant-level values were then averaged and the results are shown in Fig. 3c.
3. *<u>Headset reliability:</u>* To assess whether correlation reliability (within-visit or across-visit) depended on which headset was used, we took the following approach: (1) for both within-visit and across-visit session pairs, we computed the pairwise difference for each feature, which resulted in Δfeature_ij_ (i.e. change in a given feature between sessions i and j for all participants); (2) this quantity was then split between session pairs that used the same headset (Δfeature_ij_^SAME^) and those that use a different headset (Δfeature_ij_^DIFF^); (3) finally, an independent t-test was performed between Δfeature_ij_^SAME^ and Δfeature ^DIFF^, with p<0.05 suggesting a difference in TRR when using different headsets. The results are reported in the “Results: Reliability of resting state features” section of the manuscript.

#### Tasks

Each participant completed each task twice: once during the first visit (referred to as visit 1) and once during the second visit (referred to as visit 2). For each session, metrics of interest were first obtained at the channel level (test statistics and p-values from the GLM) as described above. The metrics described below were then considered at the module-level (resulting in one value for each of the 35 modules covering the head).

1. *<u>Dice coefficient:</u>* The Dice coefficient was computed to assess the similarity between the set of task-activated channels from visit 1, *X*, and the set from visit 2, *Y*. In order to disentangle reliable task activations and reliable scalp coupling we only considered the channels that were shared across visits for a given participant. Then for each visit, channels that were detected by a given module, *i*, and had a GLM p-value < 0.05 defined the set of significant channels for that module, yielding *X*_*i*_ and *Y_i_*. Finally, the Dice coefficient for each module was defined as 2 | *X*_*i*_ ∩ *Y*_*i*_| / (|*X*_*i*_| + |*Y*_*i*_|), or double the size of the intersection of the sets divided by the sum of the size of each set. This analysis yielded whole-head maps of Dice coefficients for each participant and each task. To assess this metric at the group-level, we averaged these maps over participants, yielding one population-average Dice coefficient map for the Auditory task (Fig. 4b) and one for the Go/No-Go task (Fig. 5b).
2. *<u>Pairwise Correlations</u>*: For each participant, visit, and module, test statistics from the GLM were averaged (where a channel was assigned to the module it was detected by). Then for each ROI (here, each module) we computed the Spearman correlation between the set of average test statistics from Visit 1 to that from Visit 2. This allowed us to not only assess the reliability of activity within regions associated with each task, but to also compare to other brain areas. The results are displayed as scatter plots for ROIs in Fig. 4c (Auditory task) and Fig. 5c (Go/No-Go task). Whole-head correlations are shown in Fig. 4d (Auditory task) and Fig. 5d (Go/No-Go task).
3. *<u>Headset reliability:</u>* This analysis was done exactly the same as the resting state with the exception that only across-visit reliability could be considered because each task was only performed once per visit.
4. *<u>Intra-class correlations:</u>* On the same module-level sets of averaged GLM test statistics from (2), we computed the ICC using the methods described in the resting state statistical analyses. Whole-head ICC results are shown in Fig. 4e (Auditory task) and Fig. 5e (Go/No-Go task).

## Acknowledgments

The authors gratefully acknowledge the software support of Gabe Lerner, Victor Szczepanski, and Eli Palmer. We would like to thank Srbouhi Uzunyan for providing support with participant scheduling and consenting and Dakota Decker for printing and installing AprilTags. We also thank the participants for their time and contribution to our study.

## Author contributions

Conceptualization: KLP, RMF. Software: JD, ZMA, EMK, SJ. Methodology: JD, EMK, ZMA, KLP. Formal analysis: EMK, ZMA, MT, NM. Investigation: MT, NM, EMK. Project Administration: MT, KLP. Data curation: NM, SJ. Visualization: EMK, ZMA, NM, MT. Writing - Original Draft: EMK, ZMA. Writing - Review and Editing: all authors. Author order was determined alphabetically. Dr. Katherine Perdue agrees to be accountable for all aspects of the work, ensuring that questions related to the accuracy or integrity of any part of the work are appropriately investigated and resolved.

## Data availability statement

The datasets used and/or analyzed during the current study are available from the corresponding author on reasonable request.

