## Supplemental Info for "Reliability of brain metrics derived from a Time-Domain Functional Near-Infrared Spectroscopy System"

### Supplementary Material

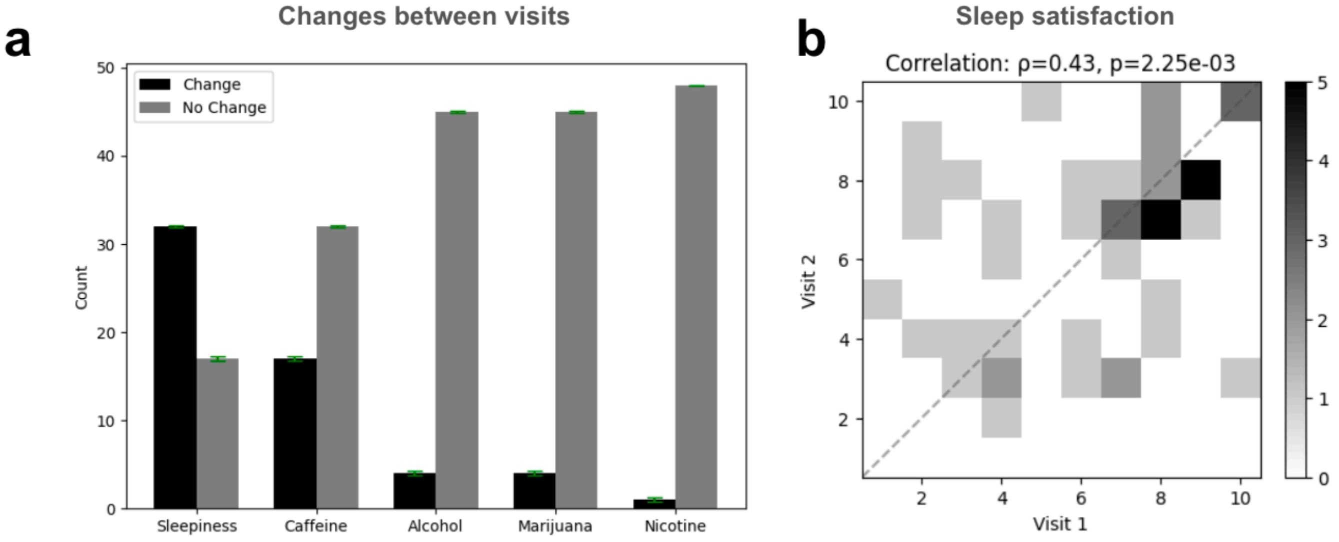


**Supplementary Figure 1. Self-reported changes in daily habits .**
**a.** Number of participants reporting a change (black bars) vs. no change (gray bars) via qualitative reports on their level of sleepiness at the time of the visit as well as recent substance use (e.g. caffeine, alcohol, marijuana, nicotine). **b.** Density plot of self-reported quantification of sleep satisfaction for visit 1 (x-axis) vs visit 2 (y-axis), with the color axis indicating the number of participants in each bin. Sleep satisfaction was on a scale of 1–10, where 10 was the highest satisfaction.

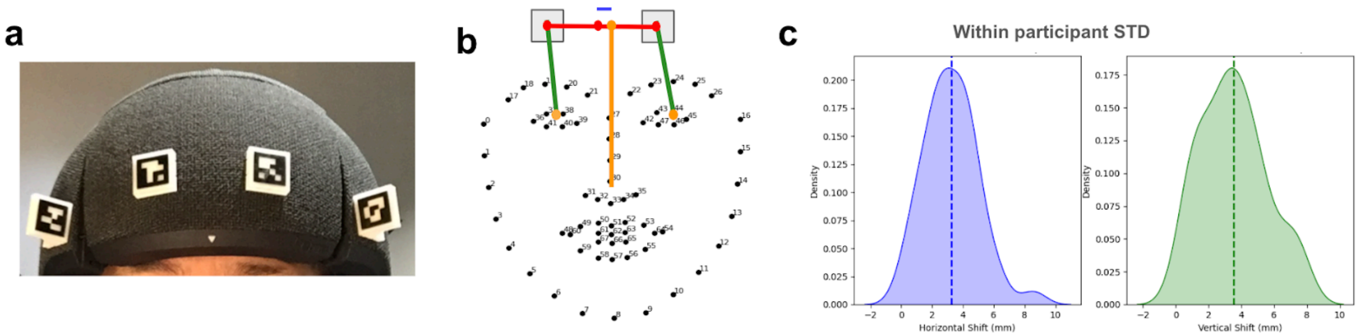


**Supplementary Figure 2. Headset placement reliability was assessed using a computer vision algorithm.**

**a.** Four distinct AprilTags were magnetically placed on each headset in fixed locations. **b.** Shown are the 68 facial landmarks from the dlib python package. Gray squares indicate the centermost AprilTags. The red dots represent the centers of the two AprilTags and the midpoint between them. The yellow indicates the points defined from the facial landmarks, centers of the eyes as well as the nose line. The green lines represent the distances between the eye and AprilTags, which were used to calculate vertical shift. The blue line between the nose line and the headset midpoint represents the horizontal shift. **c.** Kernel density estimation plots representing the within-participant variability in the horizontal and vertical shift of the headset placements. Only participants with 2 or more "good" photos were included, for a total of 42 participants. The dashed line represents the average value across participants.

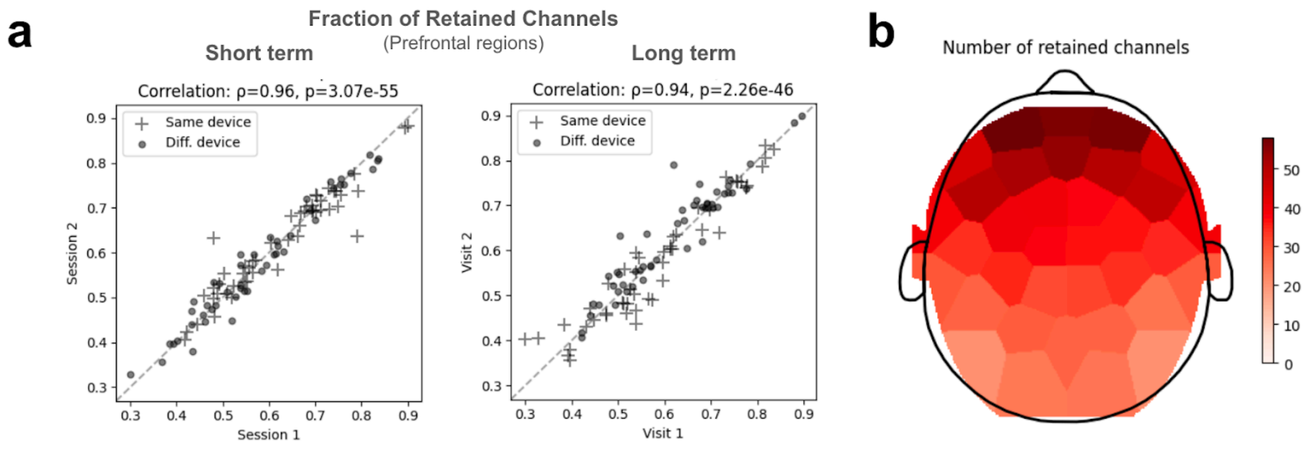


**Supplementary Figure 3. Retained channels were reliable between recording sessions.**
**a.** During resting state sessions, the fraction of retained channels in the prefrontal region was significantly and highly correlated between the sessions of a given visit (left; within-visit reliability), as well as across visits (right; across-visit reliability). The fraction of retained channels over the whole-head followed the same pattern (data not shown; Whole-head: within-visit $\rho$=0.99, across-visit $\rho$=0.98, $p<10^{-70}$ for both). **b.** Total number of retained channels (averaged across all participants) is shown for each module over the whole head (each color patch). Darker red indicates higher channel counts.

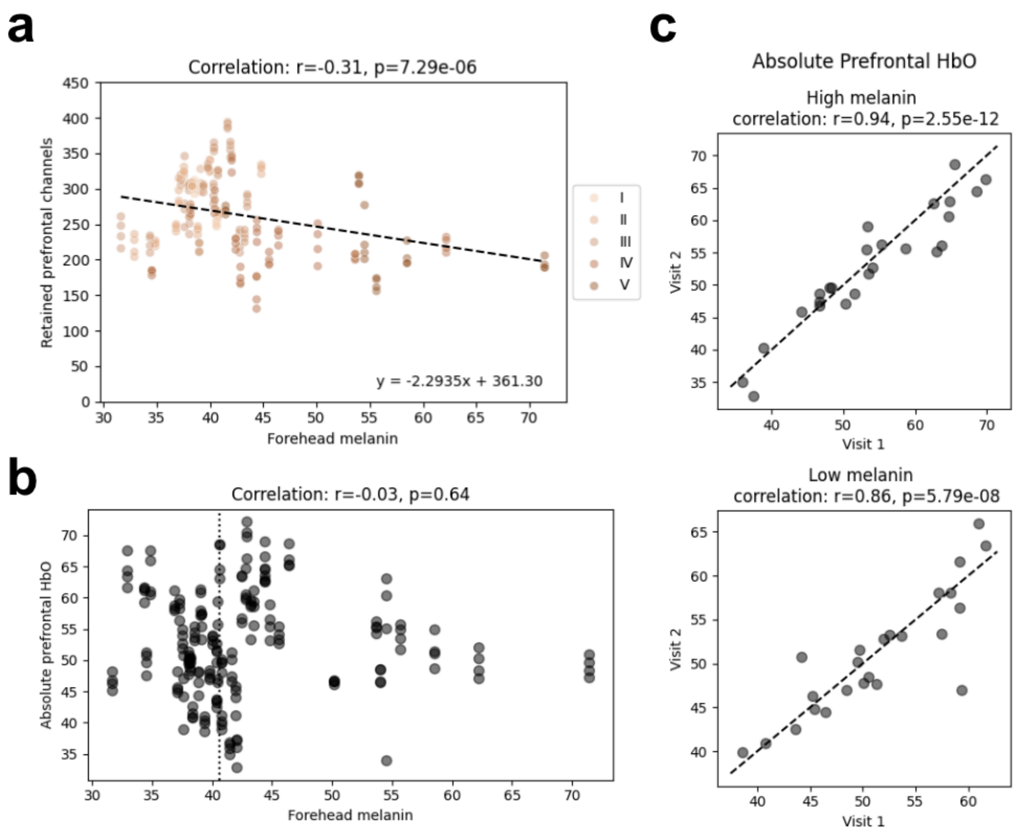


**Supplementary Figure 4. Skin color and its effect on retained channels and absolute oxygenation.**
**a.** The number of retained channels within the prefrontal region was significantly affected by participants' skin color (Melanin Index); however this effect is minimal in practical terms, as even at the highest melanin levels we can expect more than 200 prefrontal channels. Each dot is a participant and the color corresponds to the numerical values from the Fitzpatrick scale for human skin color. **b.** Absolute prefrontal HbO was not correlated with the participants' forehead melanin index (Pearson correlation). Dotted vertical line indicates the median value of the population's forehead melanin levels (median=40.61). **c.** For both low- and high-melanin groups, as defined by participants below and above the median melanin levels (40.61) respectively, absolute HbO [μM] values in the prefrontal region were significantly and highly correlated between visits. Absolute HbR, too, followed the same pattern (low melanin: r=0.93, $p=5.93x10^{-11}$; high melanin: r=0.96, $p=5.94x10^{-14}$), suggesting that the reliability in the prefrontal absolute HbO/HbR is not influenced by skin color.

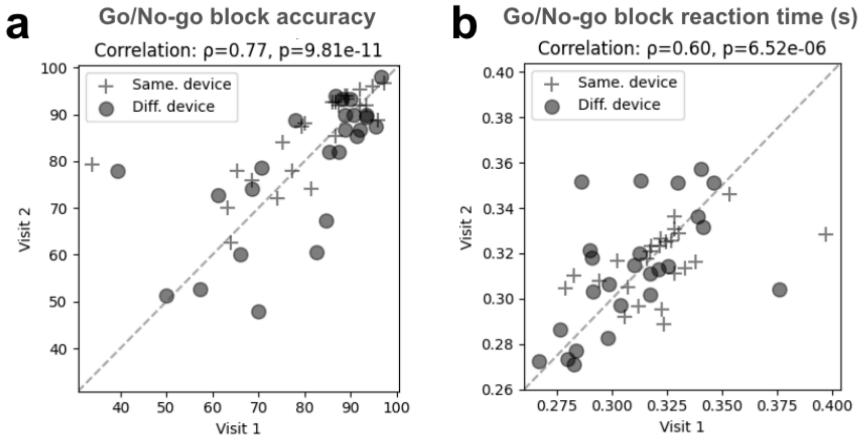


**Supplementary Figure 5. Reliability in performance during the cognitive task.**
Shown are the reliability of two different behavioral metrics during the go/no-go blocks of the cognitive task. Both accuracy (**a**) and median reaction time (**b**) showed significant correlations (Spearman) between the two visits suggesting a reliability in behavioral performance during this task.
